# Aberrant induction of p19^Arf^-mediated cellular senescence contributes to neurodevelopmental defects

**DOI:** 10.1101/2021.05.28.446099

**Authors:** Muriel Rhinn, Irene Zapata-Bodalo, Annabelle Klein, Jean-Luc Plassat, Tania Knauer-Meyer, William M. Keyes

## Abstract

Timely induction of cellular senescence is important during embryonic development and tissue repair, while aberrant activation contributes to many age-related diseases. However, whether abnormal induction of senescence in the embryo contributes to developmental defects remains unclear. To explore this, we investigated the clinically significant model of Valproic acid- (VPA) induced developmental defects. VPA is a widely prescribed drug to treat epilepsy, bipolar disorder and migraine. If taken during pregnancy however, exposure to the developing embryo can cause birth defects, cognitive impairment and Autism-Spectrum Disorder. How VPA causes these developmental defects remains unknown. We used embryonic mice and human organoids to model key features of drug exposure, including exencephaly, microcephaly and spinal defects. In the malformed tissues, in which neurogenesis is defective, we find pronounced induction of cellular senescence in the neuroepithelial cells. Critically, through genetic and functional studies, we identified p19^Arf^ as the instrumental mediator of senescence and microcephaly, but surprisingly, not exencephaly and spinal defects. Together, these findings uncover how misregulated senescence in neuroepithelial cells can contribute to developmental defects.

## Introduction

Cellular senescence is a form of permanent cell cycle arrest induced in response to a variety of stimuli. Senescence arrest is mediated by activation of cell cycle inhibitors including p21, p16^Ink4a^ and p19^Arf^ (Childs et al., 2015; Muñoz-Espín and Serrano, 2014; Rhinn et al., 2019). In addition, the arrested cells are highly secretory, producing a complex cocktail of cytokines, growth factors, extracellular matrix and other proteins, collectively known as the senescence-associated secretory phenotype (SASP). This can exert significant functional effects on the microenvironment, prominently including the activation and recruitment of immune cells to remove the senescent population. However, the SASP can also exert other effects including promoting cell proliferation, angiogenesis, epithelial-mesenchymal transition (EMT), in addition to cell plasticity and stemness (Coppé et al., 2008; Mosteiro et al., 2016; Ritschka et al., 2017). Although senescence is mostly associated with aging and disease, other studies have shown how senescent cells can have beneficial functions in various settings including embryonic development, tissue repair and regeneration, tumor suppression and reprogramming (Antelo-Iglesias et al., 2021; Demaria et al., 2014; Muñoz-Espín and Serrano, 2014; Rhinn et al., 2019). Therefore, the current view is that timely, controlled, and efficiently cleared senescent cells can have beneficial effects on tissue development and regeneration. However, when there is mis-timed or chronic induction of senescence, then this contributes to aging and disease including neurodegenerative disease, fibrosis and arthritis (Baker and Petersen, 2018; Childs et al., 2015; Rhinn et al., 2019).

During embryonic development, cells exhibiting features of senescence are detected in precise areas and at critical stages of development, including in the apical ectodermal ridge (AER) of the limb, the hindbrain roofplate, the mesonephros and the endolymphatic sac (Muñoz-Espín et al., 2013; Storer et al., 2013). Here, it is thought that the controlled induction of senescence contributes to cell fate patterning and tissue development, while the efficient removal of these cells aids in tissue remodeling (Muñoz-Espín and Serrano, 2014; Rhinn et al., 2019; Da Silva-Álvarez et al., 2019). In the embryo, senescence is mediated by p21, but appears not to involve the induction of p16^Ink4a^ and p19^Arf^, which are both expressed from the *Cdkn2a* gene (Ink4/Arf locus). Indeed, in the embryo, this locus is epigenetically silenced, and becomes active in adult life in response to oncogene expression or the aging process (Krishnamurthy et al., 2004; Liu et al., 2019; Milstone et al., 2017). Therefore, as mis-timed induction of senescence is linked with many adult diseases, we wanted to explore whether aberrant senescence might be implicated in developmental disease.

As a first model to investigate such a possible association, we investigated embryonic exposure to Valproic acid (VPA). This drug is widely used to treat a number of illnesses, including epilepsy and bipolar disorder. However, since its initial use, there have been many thousands of cases of women taking VPA during pregnancy, subsequently giving birth to children with birth defects (Clayton-Smith et al., 2019; Jentink et al., 2010; Margulis et al., 2019). In many cases, these were inadequately counseled about the associated risks, and drug use during pregnancy has continued. The main associated congenital malformations include spina bifida, facial alterations and heart malformation, with additional risk of limb defects, smaller head size (microcephaly), cleft palate and more, with higher doses associated with increased risk (Clayton-Smith et al., 2019; Jentink et al., 2010; Margulis et al., 2019). However, the most widespread consequences of VPA exposure are cognitive impairment and Autism Spectrum Disorder (ASD), which occur in 30-40% of exposed infants, and which can occur without any major physical deformity (Christensen et al., 2013; Clayton-Smith et al., 2019; Meador et al., 2013; Roullet et al., 2013).

The connection between VPA exposure and birth defects has been aided significantly by studies in rodent and primate models, leading to the hypothesis that cognitive defects arise from disruption of early neurodevelopment, around the stage of neural tube closure (Nicolini and Fahnestock, 2018; Roullet et al., 2013; Varghese et al., 2017; Zhao et al., 2019). During this period (approx. Embryonic day (E) 8.5 – E9.5 in mice), the early neuroepithelium amplifies, bends and closes to generate the neural tube, which is lined by neuroepithelial (NE) cells. During neural tube closure, the NE cells divide symmetrically to self-renew and expand (Chenn and McConnell, 1995). With the onset of neurogenesis, they differentiate into radial glial (RG) cells, that then undergo symmetric proliferative divisions to amplify their pool in the ventricular zone (VZ) of the neuroepithelium (Götz and Huttner, 2005). As development proceeds, they transition to asymmetric neurogenic divisions to produce cortical neurons directly, or indirectly by amplifying progenitors including the basal progenitors (BP) (Götz and Huttner, 2005; Miyata et al., 2001; Noctor et al., 2004). These steps must be tightly coordinated, and any perturbation of neuroepithelial or progenitor function may have consequences on later cortical neuron development which could contribute to microcephaly and other neurodevelopmental disorders including cognitive impairment and ASD.

The molecular mechanisms by which VPA perturbs development are mostly unknown, but likely result from its function as a histone-deacetylase inhibitor (HDACi) (Moldrich et al., 2013). Interestingly, in this capacity, VPA is also broadly used in cancer therapy, and has been shown to induce cellular senescence in certain settings, through direct activation of key senescence mediators including p21, p16^Ink4a^ and p19^Arf^ (Li et al., 2005). Given these associations, we investigated whether aberrant activation of senescence by VPA exposure might contribute to the associated developmental defects.

## Results

Drawing from earlier VPA-exposure studies in mice (Ehlers et al., 1992; Nicolini and Fahnestock, 2018), we established a time-course paradigm for assessing acute and developmental phenotypes caused by VPA during embryonic development (see experimental scheme Fig. 1A). Although, it has been shown that acute dosing of mice can model many key features of drug exposure in humans, mice have high drug tolerance and clearance capacity, and as such, comparatively higher doses are used to model exposure. Also, although in humans VPA causes spina bifida, a posterior neural tube closure defect where part of the spinal cord and nerves are exposed, in mice, this results in exencephaly, a defect of anterior neural tube closure where the brain is located outside of the skull (Nau et al., 1991). Here, we first analyzed E13.5 embryos from pregnant female mice that had been dosed three times around E8. As previously observed, we identified prominent and recurrent defects, such as exencephaly and spinal cord curvature (not shown) (Fig. 1B). However, we also identified that a large proportion of the mice displayed a small brain phenotype (microcephaly), a finding which was previously underestimated in mice (Fig. 1B) Next, we analyzed VPA-exposed embryos at earlier developmental stages, and could visually distinguish all phenotypes at E10.5, and even as early as E9.5, where they presented as an open neural tube and/or a smaller brain (referred to as microcephaly from here on) (Fig. 1B), and gross misalignment of the neural tube and somites (Fig. 1C). Our analyses uncover distinct separate responses to VPA that were not previously characterized, and demonstrate that VPA can cause early phenotypic changes during mouse brain development that recapitulate features of VPA-exposure in humans.

**Fig.1:**
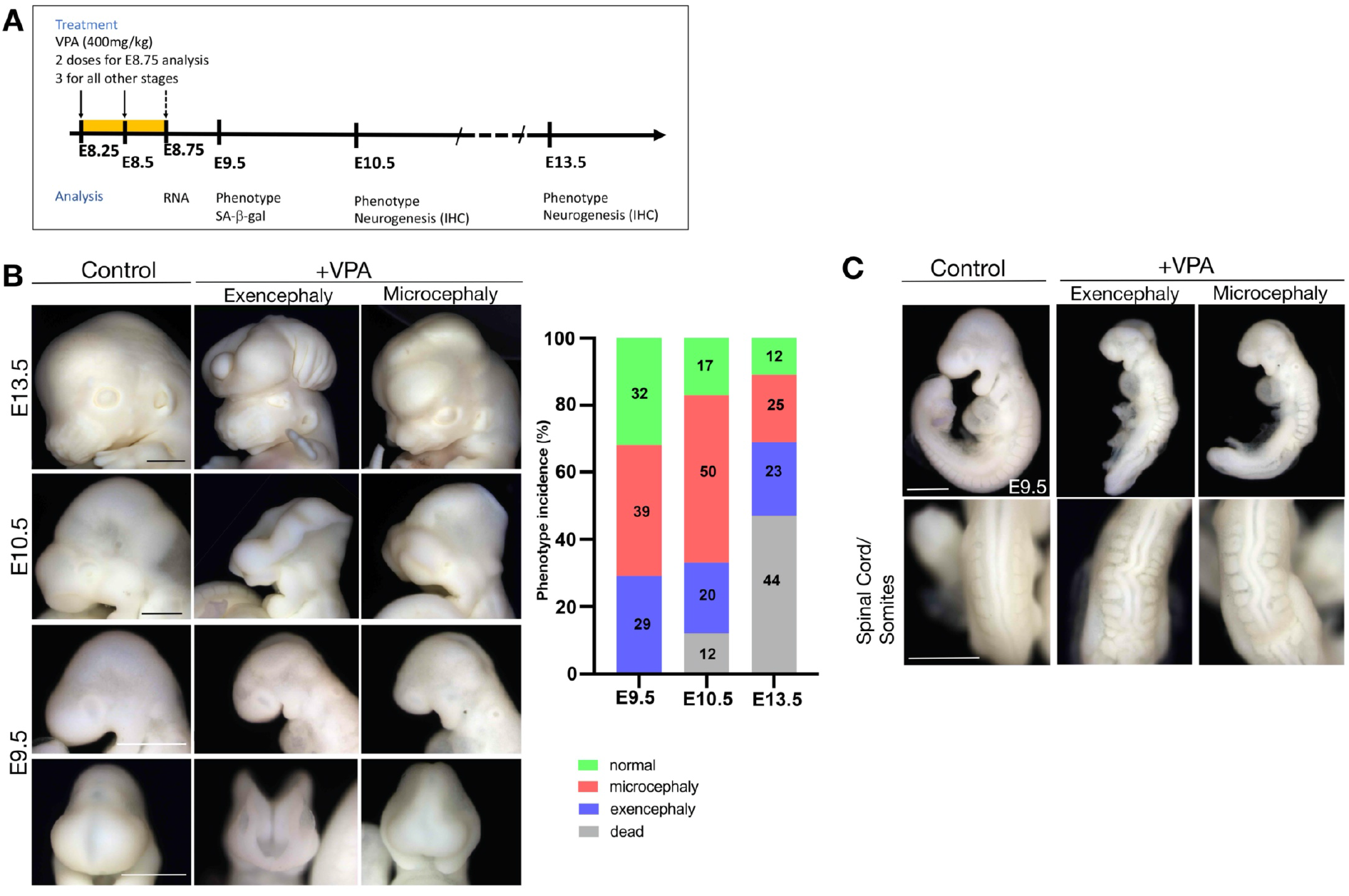
Valproic acid treatment induces developmental defects, including exencephaly, microcephaly and abnormal spinal cord development. **(A)** Schematic of experimental treatment of mice with VPA, and timeline of analysis. **(B)** Left: Embryonic head phenotypes in CD1 mice resulting from VPA exposure. Scale bar, 1mm (E13.5), 500μm (E9.5 and E10.5). Right: Phenotype incidence at different stages E9.5 (*n*= 147 embryos from 14 litters), E10.5 (*n*= 151 embryos from 16 litters) and E13.5 (*n*= 45 embryos from 4 litters). **(C)** Lateral views (top) and dorsal views (bottom) of control and VPA-treated embryos dissected at E9.5, illustrating the pronounced curve in the neural tube and abnormally shaped somites observed (*n*= 147 embryos from 14 litters). Scale bar, 500μm.

Next, we investigated whether cellular senescence was a feature in VPA-exposed mouse embryos. First, we performed wholemount staining to detect any activity of the senescence marker beta-galactosidase (SA-β-gal) on E9.5 control or VPA-exposed embryos presenting with exencephaly or microcephaly. We found that ectopic SA-β-gal activity was prominent in the forebrain and hindbrain in both exencephalic and microcephalic embryos (Fig. 2A, arrow). Unexpectedly, this ectopic staining was absent in both the spinal cord and somites. In sectioned embryos however, we found that SA-β-gal activity was prominent in the neuroepithelial cells, the embryonic precursors of neurons and glia in the brain (Fig. 2B). Upon assessing EdU incorporation, we confirmed that neuroepithelial cells were proliferative in control but not in VPA-exposed embryonic mice (Fig. 2C). In addition, while apoptotic cells were present on the surface ectoderm of the VPA-exposed embryos, as assessed by wholemount TUNEL staining, we did not detect any cell death in the neuroepithelial cells (Fig S1). We then dissected the forebrain and midbrain regions from wildtype or VPA-exposed microcephalic embryos at E8.75 and performed quantitative real-time PCR (qRT-PCR) for senescence genes, including cell cycle inhibitors and secreted components of the senescence-associated secretory phenotype (SASP). We found that *p21, p19^Arf^* and *p16^Ink4a^* and the SASP genes *IL6, IL1a, IL1b, Pai1* were strongly induced in VPA-exposed embryos (Fig 2D). Together, these data uncover that VPA induces ectopic senescence in neuroepithelial cells during developmental neurogenesis.

**Fig.2:**
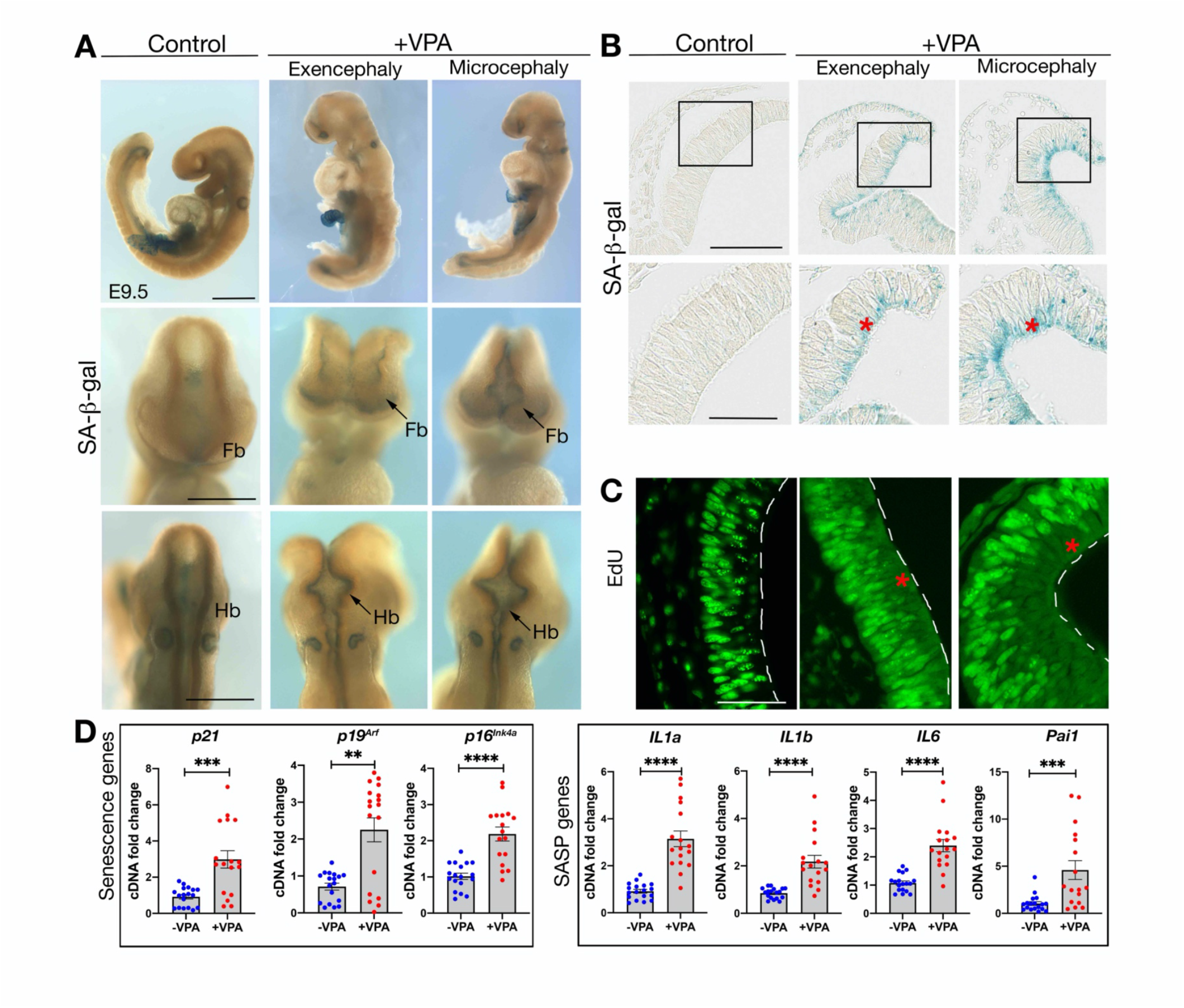
Senescence is induced in the forebrain and hindbrain neuroepithelium of VPA-treated embryos. **(A)** Whole mount SA-β-gal staining in control and VPA-treated embryos at E9.5 (*n*= 18 embryos from 7 litters). Top row, lateral views. Scale bar, 500μm. Middle row, frontal views and bottom row, dorsal views. Scale bars, 50μm. Fb, forebrain. Hb, Hindbrain. **(B)** Sections through whole mount SA-β-gal stained forebrains (Scale bar, 100μm). Box shows the region imaged in lower panel (scale bar, 50μm). Red asterisks highlight senescent cells. (*n*= 8 embryos from 4 litters). **(C)** EdU incorporation in neuroepithelial cells. Red asterisks indicate location of senescent cells (*n*= 6 embryos from 5 litters). White dashed lines indicate apical surface of the neural tube. EdU, 5-ethynyl-2’-deoxyuridine. Scale bar, 50 μm. **(D)** qRT-PCR analysis on E8.75 forebrain + midbrain, for senescence markers (*p21, p19^Arf^, p16^Ink4a^*) and SASP genes (*IL1a, IL1b, IL6* and *Pai1*) (*n*=17-18 embryos from 3 different litters). Data bars represent mean ± SEM. Mann-Whitney test: **p ≤ 0.01, ***p ≤ 0.001 and ****p ≤ 0.0001.

To investigate the potential impact of such aberrant senescence on later cortical development, we analyzed telencephalic corticogenesis at subsequent developmental stages. Neuroepithelial cells undergo differentiation into progenitors, which will then give rise to neurons and glia. When we performed immunostaining in microcephalic embryos for the neural progenitor markers Pax6 (apical progenitors), and Tbr2 (intermediate progenitors), and for the neuronal differentiation marker Tuj1, we found a significant decrease in progenitors and neurons at E10.5 in VPA-exposed embryos (Fig. S2) and E13.5. (Fig. 3). Similar results were seen in exencephalic embryos (not shown). Overall, these data associate aberrant senescence in neuroepithelial cells of the embryo with decreased neurogenesis and impaired cortical development.

**Fig.3:**
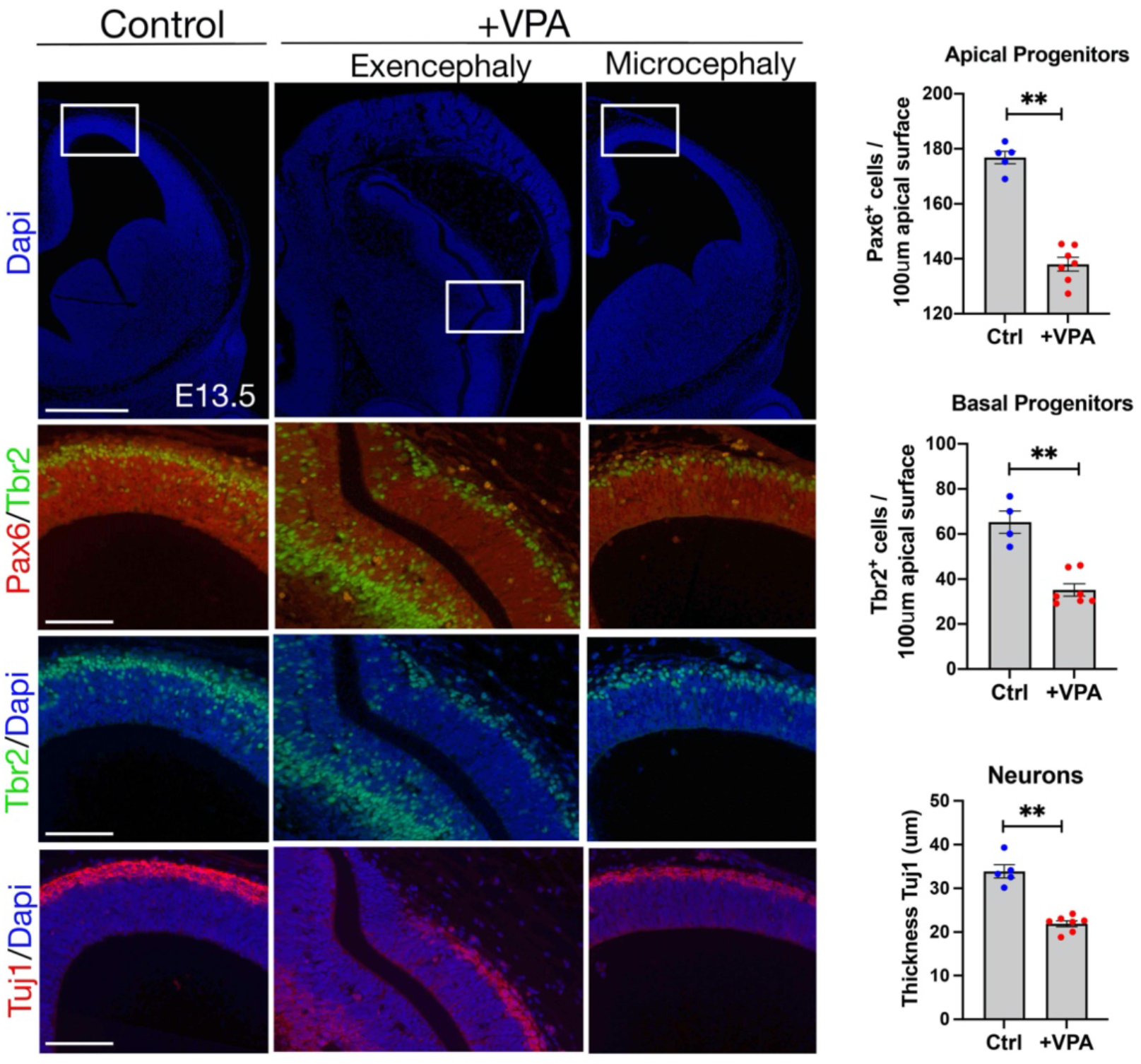
Valproic acid treatment and senescence induction is associated with decreased neurogenesis. Immunostaining for Pax6, Tbr2, Tuj1 on cortical sections (coronal) of E13.5 embryos. Box highlights the region in lower images. Scale bar, 500μm (top row), 100μm (rest). Quantification of Pax6 and Tbr2 positive progenitors or the thickness of the neuronal layer in the microcephalic cortical vesicles (for each condition, 5 embryos from at least 4 different mothers were analyzed). Data bars represent mean ± SEM. Mann-Whitney test: **p ≤ 0.01.

We next sought to assess if VPA exposure might similarly induce senescence in human neuroepithelial cells, and used cerebral organoids to investigate this possibility. We grew organoids as previously described (Lancaster et al., 2013), and exposed these to different concentrations of VPA at time points equivalent to developmental stages in mouse. Specifically, we treated cultures with 1-2mM VPA from day 18-25, and analyzed the organoids upon VPA removal at day 25, or allowed the organoids to develop until day 42, when neuronal differentiation could be assessed (Fig. 4A).

**Fig.4:**
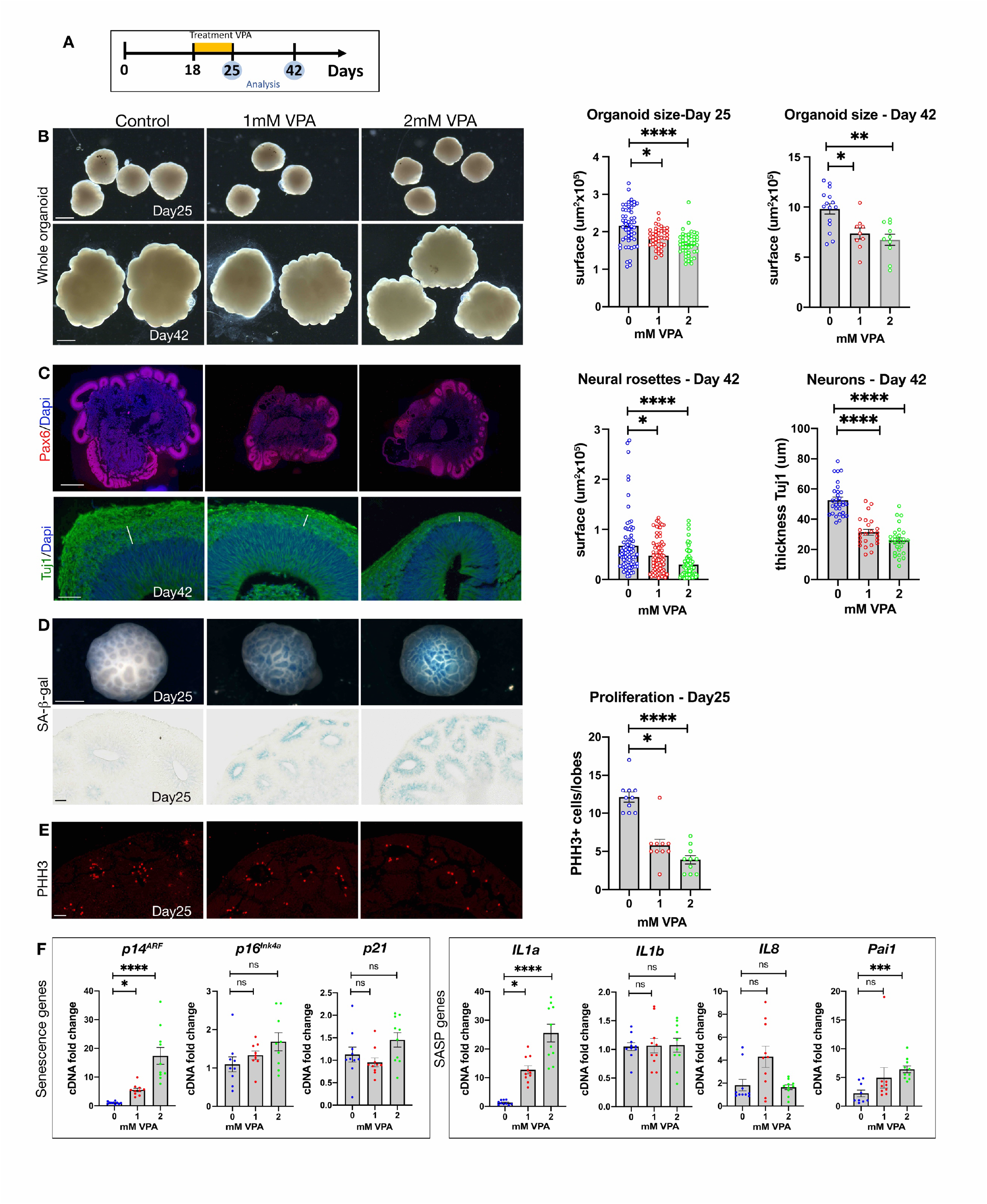
Cerebral organoids treated with VPA show a decreased size, impaired neurogenesis and induction of senescence in neuroepithelial cells. **(A)** Schematic for organoid cultures experiments. **(B)** Left: Bright field images of cerebral organoids at days 25 and day 42. Scale bar, 1 mm. Right: organoid size (μm^2^) at day 25 (*n* = 52 (Control), 41 (1mM VPA), 45 (2mM VPA), 4 independent experiments) and day 42 (*n* = 15 (Control), 9 (1mM VPA), 10 (2mM VPA), 4 independent experiments). Data bars represent mean ± SEM. Kruskal-Wallis test: *p ≤ 0.05, **p ≤ 0.01 and ****p ≤ 0.0001. **(C)** Left: Immunostaining on sections of control and VPA-treated organoids for Pax6 (red) or Tuj1 (green), counterstained with Dapi (blue). Scale bar, 500μm (Pax6) and 50μm (Tuj1). Right: Neural rosette area at day 42 (*n* = 79 (Control), 76 (1mM VPA), 79 (2mM VPA), 4 independent experiments), and neuron layer thickness (μm) at day 42 (*n* = 30 (Control), 24 (1mM VPA), 28 (2mM VPA), 4 independent experiments). Data bars represent mean ± SEM. Kruskal-Wallis test: *p ≤ 0.05, and ****p ≤ 0.0001. **(D)** Whole mount SA-β-gal staining of day 25 organoids (scale bar, 500μm). Sections show SA-β-gal staining in the neuroepithelium (scale bar, 50μm) (*n*= 5 (Control), 5 (1mM VPA), 5 (2mM VPA), 3 independent experiments). **(E)** Left: Immunostaining on sections of control and VPA-treated organoids for PHH3 (red) at day 25. Scale bar, 50μm. Right: Proliferation quantification at day 25. (n = 10 (Control), 10 (1mMVPA), 10 (2mMVPA), 2 independent experiments). Data bars represent mean ± SEM. Kruskal-Wallis test: *p ≤ 0.05 and ****p ≤ 0.0001. **(F)** qRT-PCR analysis for senescence markers (*p21, p14^ARF^, p16^INK4A^*) and for SASP genes (*IL1a, IL1b, IL8* and *Pai1*) (*n*=10 organoids from 4 independent experiments). Data bars represent mean ± SEM. Kruskal-Wallis test: ns, not significant, *p ≤ 0.05, ***p ≤ 0.001 and ****p ≤ 0.0001.

Exposure to VPA caused a significant decrease in organoid growth that persisted after drug removal (Fig. 4B). As in mice, we assessed cortical neurogenesis in VPA-treated organoids, and found a significantly reduced expression of progenitor (Pax6, Tbr2 and Sox1) and neuronal (Tuj1) markers (Fig. 4C and Fig. S3). When we assessed senescence using wholemount SA-β-gal staining, we detected a strong induction in the organoids, which upon sectioning was found to be present specifically in the neuroepithelial cells (Fig. 4D). Proliferation was also decreased in these cells, as measured by anti phospho-Histone H3 (PHH3) staining (Fig. 4E). Finally, we assessed expression of key senescence mediators by qRT-PCR at day 25. Interestingly, we observed a significant induction of *p14^ARF^* (human ortholog of *p19^Arf^*), and the SASP genes *IL1a* and *Pai1*, but no change in *p16^INK4A^* or *p21* expression (Fig. 4F).

Thus far, our experiments uncovered that exposure to VPA causes a pronounced induction of senescence in neuroepithelial cells that is associated with a marked decrease in proliferation and neurogenesis. However, we wanted to investigate if aberrant senescence is functionally coupled to the phenotypes and impaired neurogenesis. To address this, we employed genetic loss of function models deficient in the main senescence mediators *p21, p19^Arf^* or *p16^Ink4a^*. When we treated pregnant mice, each individually deficient for these genes, with VPA, and assessed E9.5 embryos, we found that *p21*- and *p16^Ink4a^* -deficient embryos had no visible improvement in phenotype and even had a slight increase in exencephaly relative to wildtype mice (Fig.5A and Fig. S4). However, although embryos deficient in *p19^Arf^* and exposed to VPA displayed no change in exencephaly or spinal curvature defects relative to wildtype mice, they were otherwise noticeably improved, as evidenced by a reduction in the incidence and/or severity of microcephaly (Fig. 5A). To validate our observations, we measured the combined forebrain and midbrain area in these embryos. At this early stage (one day after VPA exposure), we found that the forebrain/midbrain size in *p19^Arf^*-deficient embryos was significantly larger compared to wildtype VPA-exposed embryos (Fig. 5B). The lessened size reduction was also evident with *in situ* hybridization for the forebrain marker *Six3* (Fig. S5). To assess whether the size difference phenotype correlated with changes in senescence, we again assessed SA-β-gal staining, and found that VPA-exposed *p19^Arf^*-deficient mice had reduced expression in the neuroepithelial cells relative to VPA-exposed wildtype embryos (Fig. 5C). Furthermore, when assessed by qRT-PCR, *p19^Arf^*-deficiency was associated with a decrease in *p16^Ink4a^* and a reduced SASP-response (Fig. S6). In agreement with the results from human organoids, this data points to *p19^Arf^* as a mediator of VPA-induced senescence in the embryo.

**Fig.5:**
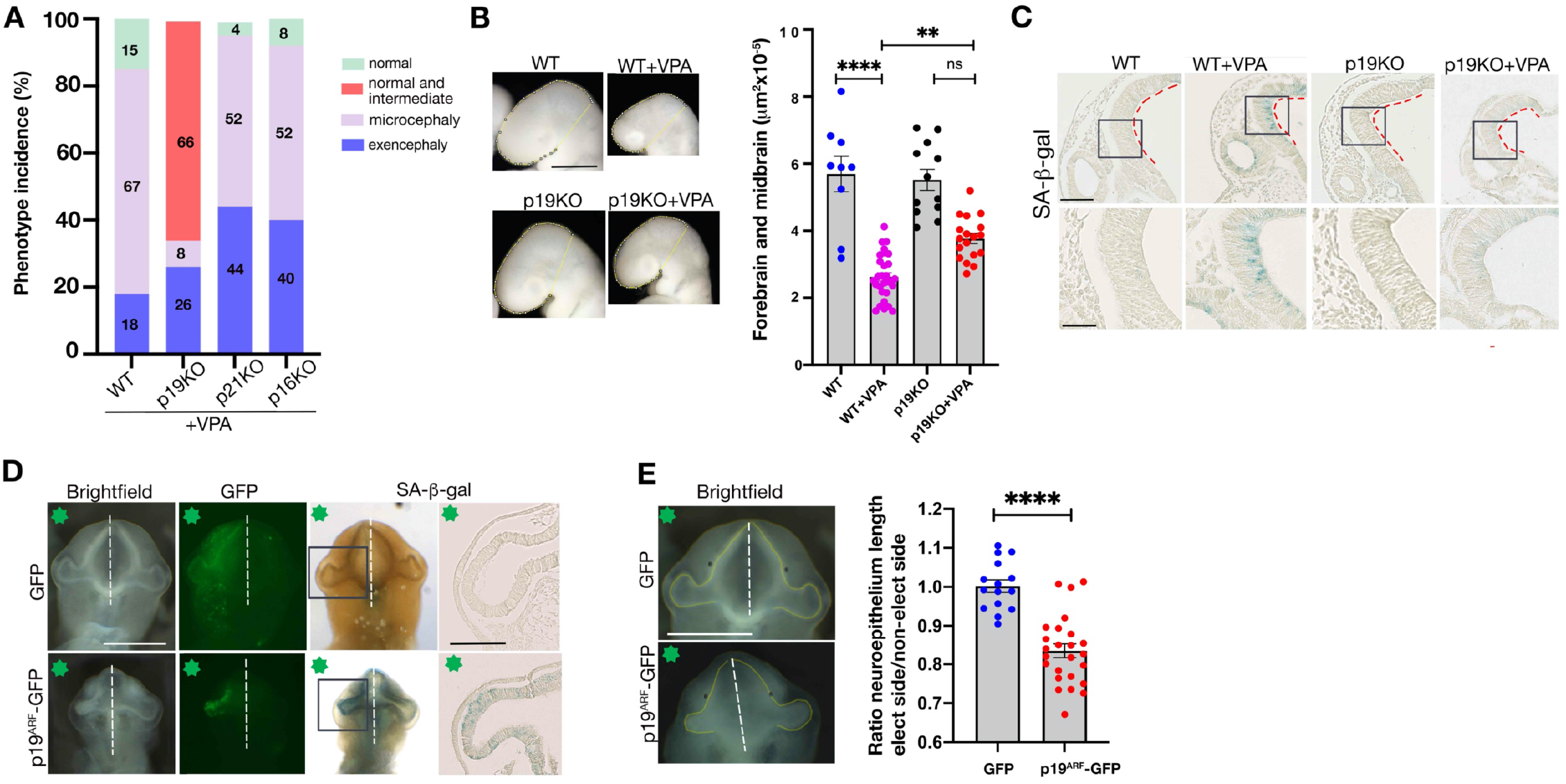
*p19^Arf^* expression causes senescence and VPA-induced microcephaly. **(A)** Phenotype incidence at E9.5 in C57/Bl6J mice. *p21*, *p19^Arf^* or *p16^Ink4a^* -deficient mice are labeled p21KO, p19KO or p16KO respectively. WT, *n*= 174 embryos from 26 litters. p19KO, *n*= 123 embryos from 19 litters, p21 KO, *n*= 72 embryos from 10 litters and p16KO, n= 97 embryos from 11 litters. **(B)** Bright field images of E9.5 embryonic heads, indicating area of the forebrain and midbrain (yellow line). Scale bar, 500μm. Graph shows surface area of forebrain and midbrain in each condition, WT, *n*= 9 embryos from 4 litters, WT+VPA, *n*= 29 embryos from 14 litters, p19KO, *n*= 12 embryos from 4 litters, p19KO + VPA, *n*= 18 embryos from 7litters. Data bars represent mean ± SEM. Kruskal-Wallis test: ns, not significant, **p ≤ 0.01 and ****p ≤ 0.0001. **(C)** Representative brain sections of E9.5 SA-β-gal stained WT or p19KO (scale bar, 100μm). Box shows the region imaged in lower panel (scale bar, 50μm). Red dashed lines indicate apical surface of the neural tube. WT, *n*= 5 embryos from 3 litters, WT+VPA, *n*= 6 embryos from 4 litters, p19KO *n*= 5 embryos from 3 litters for p19KO, p19KO + VPA *n*= 9 embryos from 3litters. **(D)** Ventral views of chick embryos at stage HH12, electroporated with a *GFP* or a *p19^Arf^-GFP* plasmid. Green star indicates electroporated side. Scale bar, 500μm. Embryos were stained for SA-β-gal activity. Boxes indicate sectioned area of forebrain neuroepithelium shown. Scale bar, 100μm. **(E)** Brightfield embryos with yellow line shows length of neuroepithelium. Scale bar, 500μm. Graph shows ratio of length of neuroepithelium in electroporated side compared to control side. GFP, *n* = 15 embryos from 4 different electroporations, *p19^Arf^-GFP, n*= 25 embryos from 9 different electroporations. Data bars represent mean ± SEM. Unpaired t-test: ****p ≤ 0.0001.

To further investigate this association and to determine if ectopic p19^Arf^ expression is sufficient to induce senescence and cause developmental defects when aberrantly expressed in the neuroepithelium, we electroporated mouse *p19^Arf^* into the neuroepithelial cells of chick embryo forebrains. In comparison to GFP-control plasmid, we found that *p19^Arf^* expression caused a unilateral perturbation of development, decreasing forebrain size, and induced strong ectopic SA-β-gal activity in the neuroepithelial cells (Fig. 5D and Fig. S7). These data demonstrate that aberrant *p19^Arf^* expression is sufficient to induce senescence and developmental defects.

Given that *p19^Arf^*-deficiency is protective for VPA-induced embryonic developmental defects, we wanted to understand the underlying mechanism at a molecular level. To this end, we performed RNA-sequencing on the forebrain/midbrain region from both wildtype and *p19^Arf^*-deficient embryos, either treated or untreated with VPA. Through phenotype pathway analysis of differentially expressed genes it was evident that many neurodevelopmental and ASD-related phenotypes, including exencephaly and microcephaly, were associated with significantly downregulated genes in VPA-exposed wildtype mice. Specifically, these gene signatures were associated with Wnt and Hippo signaling (Kwan et al., 2016) (Fig. 6A and Fig. S8A). In *p19^Arf^*-deficient animals however, most of these signatures were significantly less affected, confirming our phenotypic observations of the genetic backgrounds (Fig. 6A and Fig. S8A).

**Fig.6:**
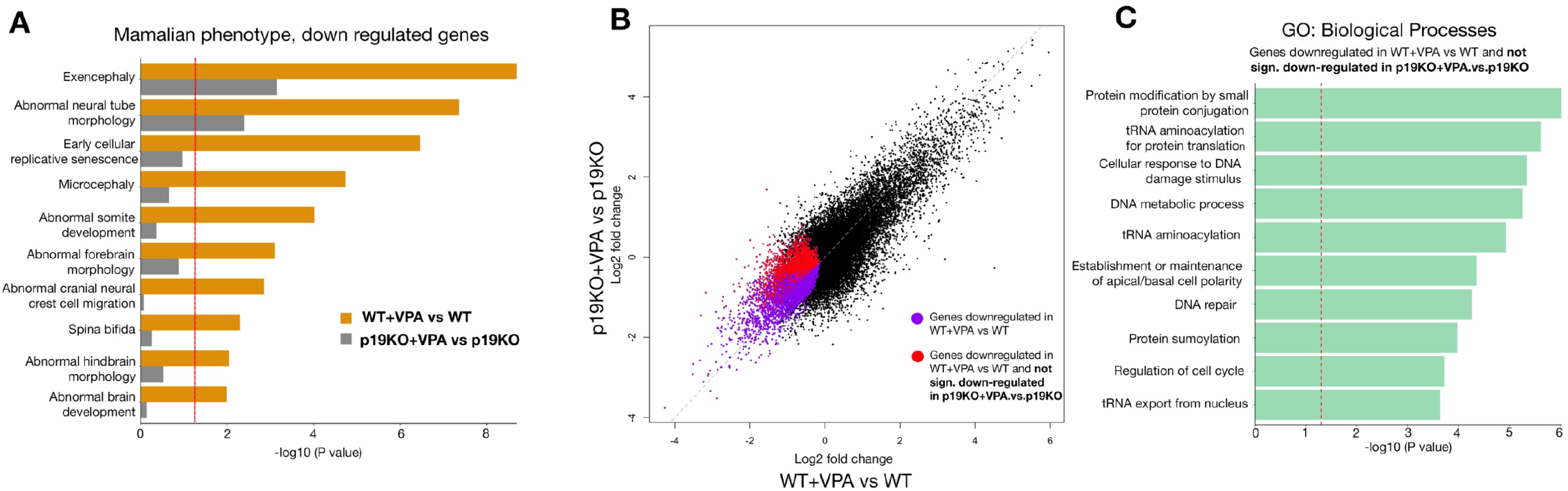
*p19^Arf^* deficiency rescues VPA-induced gene signatures associated with neurodevelopmental defects. **(A)** Mammalian Phenotype pathway analysis on the downregulated genes from RNA-seq of the forebrain and midbrain **(B)** Scatter plot showing mRNA fold changes for the genes in WT+VPA compared to WT, and in p19KO + VPA compared to p19KO. **(C)** GO Biological process analysis on genes highlighted in E with red dot.

Genetic population studies have identified candidate genes associated with microcephaly and ASD (Jayaraman et al., 2018; Takata et al., 2018). Many of these genes are significantly decreased in both the forebrain and midbrain of VPA-exposed wildtype embryos, including Chd8, Dyrk1a, Fmr1, Cep63, and others. However, most were not restored upon *p19^Arf^*-loss (Fig. S8 B,C), suggesting that senescence may be regulated independently or downstream of these specific genes. Therefore, to get a better understanding of how p19^Arf^ might induce these ectopic phenotypes, we analyzed the subset of genes that were significantly downregulated in VPA-exposed wildtype embryos, but that were not significantly decreased in *p19^Arf^*-deficient embryos (red genes in Fig 6B). Within this p19^Arf^-dependent gene set, we identified tRNA aminoacylation and tRNA export (Fig. 6C and Fig. S8D). Interestingly, perturbation of tRNAs or their regulatory mechanisms is linked to microcephaly and neurodevelopmental defects (Schaffer et al., 2019). This suggests that p19^Arf^-mediated senescence and repression of these genes may contribute to microcephaly and cognitive impairment.

Finally, to conclusively demonstrate that aberrant senescence contributes to impaired neurodevelopment, we asked whether p19^Arf^ deficiency would rescue some of the major defects caused by VPA exposure. To answer this question, we measured progenitor and neuronal status during cortical neurogenesis at later stages, when neurodevelopment has progressed further. As before, wildtype embryos exposed to VPA and examined at E13.5 presented with a significant reduction in the number of progenitors and neurons (Fig. 7). Strikingly however, *p19^Arf^*-deficient mice were not as susceptible to VPA exposure, and presented with significantly increased numbers of progenitors, as well as increased thickness of the neuronal zone relative to wildtype embryos. These experiments conclusively demonstrate that p19^Arf^, in response to VPA, drives a senescence-mediated block in neurogenesis.

**Fig.7:**
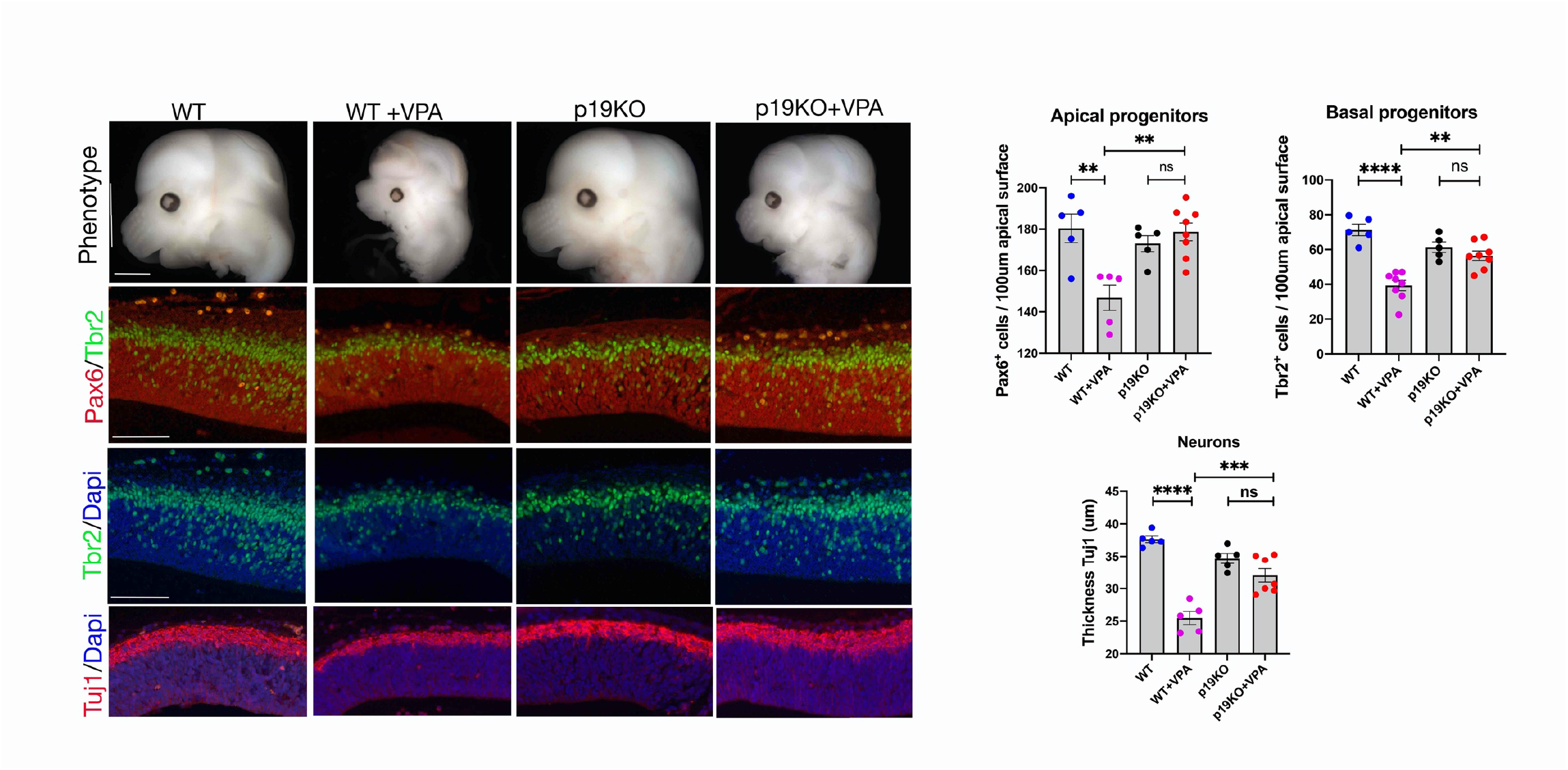
*p19^Arf^* deficiency rescues VPA-induced microcephaly. Images showing the cortical vesicles. Scale bar, 1mm. Immunostaining on cortical sections, E.13.5, for Pax6, Tbr2, Tuj1 and counterstained with Dapi. Scale bar, 100μm. Graphs showing number of Pax6 and Tbr2 positive progenitors or the thickness of the Tuj1 neuronal layer in the cortical vesicles (for each condition, minimum 5 embryos from at least 4 different mothers were analyzed). Data bars represent mean ± SEM. One-way ANOVA plus Tukey post-hoc test: ns, no significant, **p ≤ 0.01, ***p ≤ 0.001 and ****p ≤ 0.0001.

## Discussion

Together, these findings demonstrate that aberrantly induced senescence perturbs embryonic development, leading to developmental defects, and also advances our understanding of how VPA causes neurodevelopmental disorders.

A major finding of this work is that it makes an exciting functional connection between aberrant cellular senescence and developmental defects. While abnormal induction or chronic accumulation of senescent cells has been linked to many adult and age-related diseases, we demonstrate here a causative role for senescence in neurodevelopmental defects. Interestingly, we identify that the neuroepithelial cells are the site of senescence induction. As this population of cells is a critical precursor of all mature cell types in the brain, it stands to reason that this is one of the most perturbed population of cells in neurodevelopmental disorders. We demonstrate that induction of senescence in the NE cells correlates with a subsequent impairment in corticogenesis and neural differentiation, that is rescued in the absence of a key senescence gene. This demonstrates that this induction of senescence effectively blocks the development of the affected neuroepithelial cells. As the majority of infants with problems associated with VPA exposure have cognitive defects, including developmental delay and ASD, this suggests that senescence in the NE cells could be a significant contributor to these outcomes.

This study also links aberrant senescence in the NE cells with microcephaly. Indeed, microcephaly is a feature of VPA-exposure in infants, and the strategy used here in mice of an acute model of VPA exposure, mimics many associated features of VPA exposure in humans (Clayton-Smith et al., 2019; Nicolini and Fahnestock, 2018). Such high-dose acute treatment is necessary to avoid the low penetrance of developmental defects seen in mice. Of course, it is possible that this may exaggerate some of the features found in humans. However, as shown by VPA treatment of human organoids during week-long exposure, the outcome of senescence in NE cells is conserved, correlating with increased expression of *p14^Arf^* and decreased neurogenesis. As affected human embryos are chronically exposed to the drug during development, it is possible that this would cause a lower, but longer incidence of senescence in NE cells or their derivatives, but which may perturb differentiation in specific areas or at different stages of development, yet without always manifesting as microcephaly. Interestingly however, there is a strong correlation between microcephaly at birth and lower cognitive ability in ASD patients (Courchesne et al., 2003, 2007; Fombonne et al., 1999; Mraz et al., 2007; Zachor and Ben-Itzchak, 2016), suggesting that further exploration for possible connections between mistimed senescence during development and ASD is warranted.

How VPA causes birth defects has remained unclear, but exposure during the first trimester, around the stages of neural tube closure is suggested as being critical in driving the phenotypes associated with this drug, and with higher doses associated with increased risk (Clayton-Smith et al., 2019; Roullet et al., 2013; Zhao et al., 2019). Our findings identify that the drug can affect individual embryos differently, causing severe physical defects such as exencephaly in some, while causing different effects such as microcephaly in others. The reasons for this varied response remain unknown, but is likely related to cell-type specific responses. For example, while we did not detect aberrant cell death in the NE cells, apoptosis was apparent on the surface ectoderm after VPA-treatment, suggesting that in some cases, VPA-induced cell death could prevent neural tube closure, causing exencephaly.

An outstanding question as to why there is such a restricted pattern of senescence induced in the embryo by VPA, is likely related to the pattern of expression of HDAC genes. Valproic acid is an HDACi, interfering in particular with HDACs 1 and 2. HDACs have distinct patterns of expression in the embryo, with HDAC 1 and 2 being prominent in the early brain, thereby likely making cells in this region susceptible to effects of the drug (D’Mello, 2020; Jaworska et al., 2015; Murko et al., 2010). The *Ink4/Arf* locus is directly repressed by HDACs, which contributes to the normal silencing of these genes in the embryo (Milstone et al., 2017). However, HDACi’s and VPA can directly de-repress this locus, in particular *p19^Arf^* (Matheu et al., 2005; Soriano-Cantón et al., 2015). Importantly, we demonstrate that ectopic induction of *p19^Arf^* is sufficient and able to cause ectopic senescence, impaired neurogenesis and developmental defects.

It might also be considered surprising that the senescence-induced phenotypes are mediated by p19^Arf^, and not p16^Ink4a^, the latter of which is often considered a primary mediator of adult and age-associated senescence (Liu et al., 2019; Muñoz-Espín and Serrano, 2014; Rhinn et al., 2019). One possibility may relate to the timing and duration of senescence. Interestingly, in senescence induced in cells in culture, p16^Ink4a^ expression often appears later in the program. Perhaps in this case in the embryo, the senescent cells are transiently induced following VPA-exposure, and are ultimately cleared before expression of p16^Ink4a^ can manifest. In addition, it does appear that this specific inducer, VPA, has preferential ability for activating p19^Arf^ over p16^Ink4a^ (Matheu et al., 2005; Soriano-Cantón et al., 2015). Therefore, it is possible that p16^Ink4a^, or even mis-expressed p21 could contribute to other developmental defects.

Furthermore, although VPA is an HDACi, which are typically associated with gene-activation, we find, as did others, that the developmental phenotypes are associated with the down-regulated and not the up-regulated genes (Takata et al., 2018). This suggests that VPA induction of p19^Arf^-mediated senescence causes a broad repression of key developmental pathways, which impact NE fate and contribute to the developmental phenotypes, as many of these were rescued in the absence of *p19^Arf^*. Among these, we identify tRNA regulation as one of the most significantly restored pathways in the absence of *p19^Arf^*. Importantly, p19^Arf^ can directly block tRNA synthesis (Morton et al., 2007), while disruption of tRNA function is strongly associated with microcephaly and neurodevelopmental disorders (Blanco et al., 2014; Hoye and Silver, 2021; Kuo et al., 2019; Schaffer et al., 2019; Wang et al., 2020; Zhang et al., 2014). Interestingly, recent findings also show that induction of senescence involves disruption of tRNA expression, further reinforcing this link (Guillon et al., 2020). It will be interesting to determine whether such inhibition of tRNA function contributes to specific, or global alterations in protein translation in senescent cells, either in VPA-induced developmental defects, or other settings.

Overall, the discovery that atypical activation of senescence in the embryo can perturb development raises the intriguing possibility that it may also contribute to defects in developmental contexts beyond those we studied here, and highlights how the study of mistimed senescence in developmental disorders merits further study.

## Materials and Methods

### Animal maintenance and VPA administration

Pregnant CD1, C57Bl6/J, *p21-/-, p19^Arf^-/-* and *p16^Ink4a^ -/-* were maintained in a temperature- and humidity-controlled animal facility with a 12h light/dark cycle. We administrated 400mg/kg (Sigma-aldrich) VPA or PBS as control, intra-peritoneally to timed-pregnant females, at embryonic day (ED) 8 (3 times (9am, 1pm, 4pm). The *p21-/-, p19^Arf^-/-* and *p16^Ink4a^ -/-* mice were on a C57BL6J background, so were compared to C57BL6J wildtype as control. We observed that the C57Bl6J mice are more sensitive than the CD1 mice to induction of microcephaly (Fig. 1A and Fig. 4A). For qRT-PCR and RNA-seq analysis, only the first 2 doses were administered, and samples were collected at E8.75. All the experimental procedures were in full compliance with the institutional guidelines of the accredited IGBMC/ICS animal house, in compliance with French and EU regulations on the use of laboratory animals for research, under the supervision of Dr. Bill Keyes who holds animal experimentation authorizations from the French Ministry of Agriculture and Fisheries (#12840).

### Organoids

Cerebral organoids were generated from the iPSC line HPSI0214i-kucg_2 (Catalog# 77650065, HipSci) using the STEMdiff™ Cerebral Organoid Kit (Catalog #08570 and #08571) from StemCell Technologies. Representative pictures were acquired with a LEICA DMS 1000. We acknowledge Wellcome Trust Sanger Institute as the source of HPSI0214i-kucg_2 human induced pluripotent cell line which was generated under the Human Induced Pluripotent Stem Cell Initiative funded by a grant from the Wellcome Trust and Medical Research Council, supported by the Wellcome Trust (WT098051) and the NIHR/Wellcome Trust Clinical Research Facility, and acknowledges Life Science Technologies Corporation as the provider of “Cytotune.”

### Immunofluorescence

Embryos and organoids were fixed in 4% PFA for 30 min at 4°C, washed in PBS, and processed for paraffin embedding. Sections were obtained using a microtome (8 μm, Leica 2035 Biocut). After antigen unmasking in citrate buffer (0.01 M, pH 6) for 15 min in a microwave oven, slides were blocked with 5% donkey serum, 0.1% TritonX-100 in phosphate-buffered saline (PBS) and incubated overnight with the following primary antibodies: phospho-histone H3 (1:500, Upstate #05-806); Pax6 (1:300, Covance #PRB-278P); Tbr2 (1:300, eBioscience #14-4875); Sox1 (1:50, R&D systems #AF3369); βIII-tubulin/Tuj1 (1:200, Covance #MMS-435P-100); p19^Arf^ (5-C3-1) rat monoclonal antibody (Santa-Cruz #sc-32748); GFP 2A3 (IGBMC). Primary antibodies were visualized by immunofluorescence using secondary antibodies from donkey (1:400, Invitrogen: Alexa Fluor 568 donkey anti-mouse IgG #A-100037, Alexa Fluor 488 donkey anti-rat IgG #A-21208, Alexa Fluor 488 donkey anti-rabbit #A-21206, Alexa Fluor 568 donkey anti-Goat IgG #A-11057) and from goat (1:400, Invitrogen: Alexa Fluor 568 goat anti-rabbit IgG #A-110111,Alexa Fluor 488 goat anti-mouse IgG #A11001, Alexa Fluor 568 anti-rat IgG #A11077), and cell nuclei were identified using DAPI (1:2000). Stained sections were digitized using a slide scanner (Nanozoomer 2.0-HT, Hamamatsu, Japan), and measurements (thickness of the neuronal layer) were performed using the NDPview software of the digital scanner.

### SA-ß-gal staining

Whole-mount SA-ß-gal, was detected as previously described (Storer et al., 2013). Incubation with X-gal was performed overnight for mouse embryos and 1h30 for organoids. For determination of specific localization of senescence in embryonic tissue, embryos stained with SA-ß-gal were post-fixed in 4% PFA overnight at 4°C, embedded in paraffin and sectioned. Representative pictures were acquired using a macroscope (Leica M420) and stained sections were digitized using a slide scanner (Nanozoomer 2.0-HT, Hamamatsu, Japan)

### EdU

To assess cell proliferation in embryos, pregnant female mice at E9.5 were injected intraperitoneally with 5-ethynyl-2’-deoxyuridine (EdU; 50 mg/kg body weight) for 1 hr. Click-iT® EdU Alexa Fluor® 488 Imaging Kit (Thermofisher) was used as per manufacturers protocol. Representative pictures were acquired using a microscope (DM4000B).

### TUNEL

Cell death was assessed using the TdT-mediated dUTP nick end-labeling (TUNEL) method (ApopTagPeroxidase In Situ Apoptosis detection kit, Millipore) as per manufacturers’ instructions. Representative pictures were acquired using a macroscope (Leica M420) and a microscope (DM4000B).

### RT-qPCR and analysis

The combined forebrain and midbrain region was manually dissected from E8.75 embryos, and snap-frozen. RNA was extracted from individual embryos using the RNAeasy mini kit (Qiagen). 10 ng RNA were used for analysis with the LUNA one-step RT-qPCR kit (LUNA E3005L Biolabs). The relative expression levels of the mRNA of interest were determined by real-time PCR using Quantifast SYBR Green Mix (Qiagen) with specific primers listed in Supplementary Table 1 and a LightCycler 480 (Roche). Samples were run in triplicate and gene of interest expression was normalized to human Gapdh or mouse Rplp0.

### In ovo electroporation

Fertilized chicken embryos were obtained from local farmers. Chick eggs were incubated in a humidified chamber at 37°C, and embryos were staged according to (Hamburger and Hamilton, 1951). 1.5 μg/μL DNA constructs (*pCAGGS-GFP* [a gift from Dr J. Godin, IGBMC] or *pCAGGS-p19^ARF^-GFP* [*p19^Arf^* coding sequence was cloned in XhoI/NheI multiple cloning sites in the *pCAGGS-GFP]*) mixed with 0.05% Fast Green (Sigma) were injected into neural tubes of stage HH8 chick embryos and electroporated on the right side, leaving the left side as untreated control. Electroporation was performed using a square wave electroporator (BTX ECM 830 electroporation system) and the parameters applied: three pulses of 15V for 30ms with an interval of 1s. Embryos were harvested 24 hours after electroporation and processed for SA-ß-gal, histology and immunohistochemistry. Representative pictures were acquired using a macroscope (Leica Z16 APO) and a microscope (Leica DM4000B).

### Whole-mount in situ hybridization

RNA probes were prepared by *in vitro* transcription using the Digoxigenin-RNA labeling mix (Roche). Template plasmids were kindly provided by Drs G. Oliver (*Six3*) and S.L. Ang (*Mox1*). Mouse embryos were dissected in ice-cold PBS and fixed O/N in 4% paraformaldehyde (PFA)/PBS. After several washes in PBS1X/0.1% Tween-20 (PBT), embryos were bleached for 1 h in 3% H_2_O_2_/PBT and washed in PBT before being digested with Proteinase K (10mg/ml) for 2min. Digestion was stopped by 5 min incubation in 2 mg/ml glycine/ PBT. Embryos were washed again in PBT before post-fixing for 20 min in 0.2% glutaraldehyde/4% PFA/PBS. After further washes they were incubated in prewarmed hybridization buffer (50% deionized formamide, 5XSSC, 1%SDS, 100μg/ml tRNA) and prehybridized for 2 h at 65 °C. The buffer was then replaced with fresh prewarmed hybridization buffer containing the digoxigenin labeled RNA probes and incubated O/N at 65 °C. The next day, embryos were washed twice in buffer 1 (50% formamide; 5XSSC; 1%SDS) at 65 °C then in buffer 2 (NaCl 500mM, 10mM TrisHCl pH=7.5, 0.1%Tween20) at room temperature before treating them with RNaseA (100mg/ml) to reduce background. The embryos were rinsed in buffer 2, then in buffer 3 (50% formamide, 2XSSC). Finally, the embryos are rinsed in TBS/0.1% Tween-20 (TBST) then blocked for 2 h in 2% blocking solution (Roche) and incubated O/N in the same solution containing 1:2,500 anti-digoxigenin antibody (Roche). The next day the embryos were washed in TBST, before washing them in NTMT (NaCl 100mM, Tris-HCl 100mM pH=9,5, MgCl_2_ 50mM, Tween20 at 0.1%) and developing the signal in the dark with staining solution (4.5 μl/ml NBT and 3.5 μl/ml BCIP (Roche) in NTMT buffer).

### RNA sequencing

RNA was collected as for qRT-PCR. Full length cDNA was generated from 10 ng of total RNA from 4 individual embryos per treatment, using Clontech SMART-Seq v4 Ultra Low Input RNA kit for Sequencing (Takara Bio Europe, Saint Germain en Laye, France) according to manufacturer’s instructions with 8 cycles of PCR for cDNA amplification by Seq-Amp polymerase. Six hundred pg of pre-amplified cDNA were then used as input for Tn5 transposon tagmentation by the Nextera XT DNA Library Preparation Kit (96 samples) (Illumina, San Diego, CA) followed by 12 cycles of library amplification. Following purification with Agencourt AMPure XP beads (Beckman-Coulter, Villepinte, France), the size and concentration of libraries were assessed by capillary electrophoresis using the Agilent 2100 Bioanalyzer.

Sequencing was performed on an Illumina HiSeq 4000 in a 1×50bp single end format. Reads were preprocessed using cutadapt 1.10 in order to remove adaptors and low-quality sequences, and reads shorter than 40 bp were removed from further analysis. Remaining reads were mapped to Homo sapiens rRNA sequences using bowtie 2.2.8, and reads mapped to those sequences were removed from further analysis. Remaining reads were aligned to mm10 assembly of Mus musculus with STAR 2.5.3a. Gene quantification was performed with htseq-count 0.6.1p1, using “union” mode and Ensembl 101 annotations. Differential gene expression analysis was performed using DESeq2 1.16.1 Bioconductor R package on previously obtained counts (with default options). P-values were adjusted for multiple testing using the Benjamini and Hochberg method. Adjusted p-value <0.05 was taken as statistically significant.

Pathway analysis was performed using Enrichr (http://amp.pharm.mssm.edu/Enrichr.) with Gene Ontology 2018 and MGI Mammalian Phenotype Level 4 2019 databases. Adjusted p-value of <0.25 was used as a threshold to select the significant enrichment. The sequencing data have been uploaded to the Gene Expression Omnibus (GEO) database with accession number pending.

### Counting and Statistical analysis

For cell number quantification, positive cells for a given marker (Pax6, Tbr2 or Tuj1) were counted in a 100μm width columnar area from the ventricular zone to the apical surface in similar regions in the cortex. Immunofluorescence analyses, area measurements, and RNA expression were statistically analyzed using Prism (GraphPad, San Diego, California, United States). At least five animals of each treatment from three different litters were analyzed. Cell counting was performed on three adjacent sections. Results are presented as mean ± S.E.M. Statistical analysis was carried out employing the Mann-Whitney test for unpaired variables. For 3 or more groups, normal multiple comparisons were tested with one-way ANOVA plus Tukey post-hoc test and non-normal multiple comparisons were tested using Kruskal–Wallis test followed by a Dunn’s test. P-values < 0.05 were considered significant ( *p ≤ 0.05, **p ≤ 0.01, ***p ≤ 0.001 and ****p ≤ 0.0001).

## Acknowledgements

We thank Travis Stracker, Juliette Godin, Michele Studer, Pura Munoz, Birgit Ritschka, Christina Lilliehook (Life Science Editors) and members of the Keyes lab for comments on the manuscript. We thank the core facilities at the IGBMC for excellent technical support, including the Sequencing, Cell Culture and Microscopy platforms, the mouse facilities of the IGBMC and the Mouse Clinical Institute (ICS), and in particular Sylvie Falcone, Amélie Freismuth, Marion Humbert, Jean-Marie Garnier and Olivia Wendling for technical assistance. Sequencing was performed by the GenomEast platform, a member of the “France Génomique” consortium (ANR-10-INBS-0009). Work in the Keyes lab was funded in part by grants from: La Fondation Recherche Medicale (FRM) (AJE20160635985), Fondation ARC (PJA20181208104), IDEX Attractivité - University of Strasbourg (IDEX2017), La Fondation Schlumberger pour l’Education et la Recherche FSER 19 (Year 2018)/FRM, ANR (ANR-19-CE13-0023-03) and La Ligue Contre le Cancer. Work was also supported by the grant ANR-10-LABX-0030-INRT, a French State fund managed by the Agence Nationale de la Recherche under the frame program Investissements d’Avenir ANR-10-IDEX-0002-02.

---

**Figure S1.**
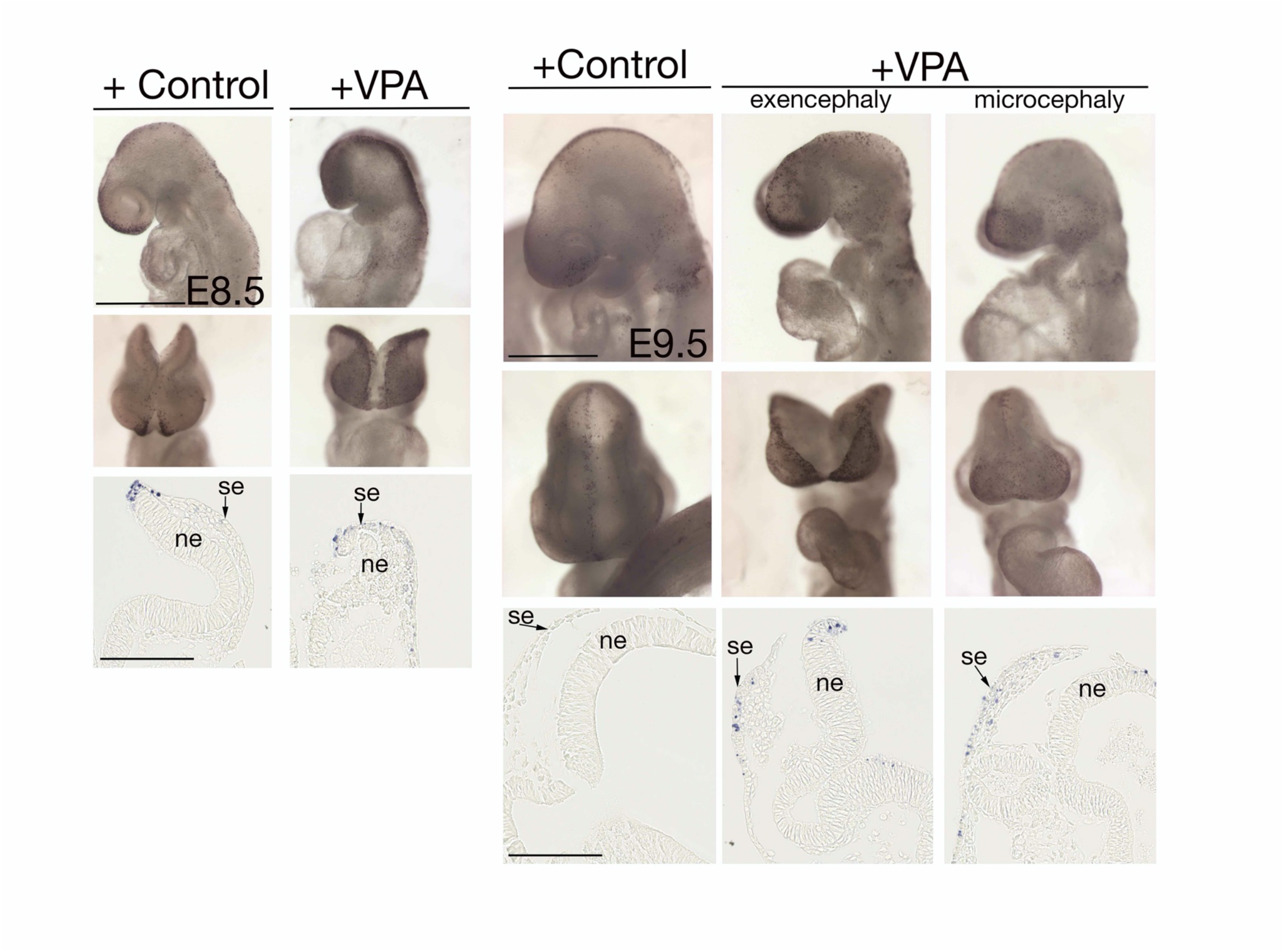
VPA does not induce apoptosis in the forebrain neuroepithelium. Control and VPA-treated embryos were stained with whole mount TUNEL assay, to assess cell death. (Left) Lateral views and frontal views of control and VPA-treated embryos dissected at E8.5. Scale bar, 500μm. Corresponding horizontal sections at the forebrain level (3 embryos from at least 2 litters were analyzed). Scale bar, 100μm. (Right) Lateral views and frontal views of control and VPA-treated embryos dissected at E9.5 (6 embryos from at least 5 litters were analyzed). Scale bar, 500μm. Corresponding horizontal sections at the forebrain level. Scale bar, 100μm. Some apoptotic cells are observed in the surface ectoderm. ne = neuroepithelium, se = surface ectoderm.

**Figure S2.**
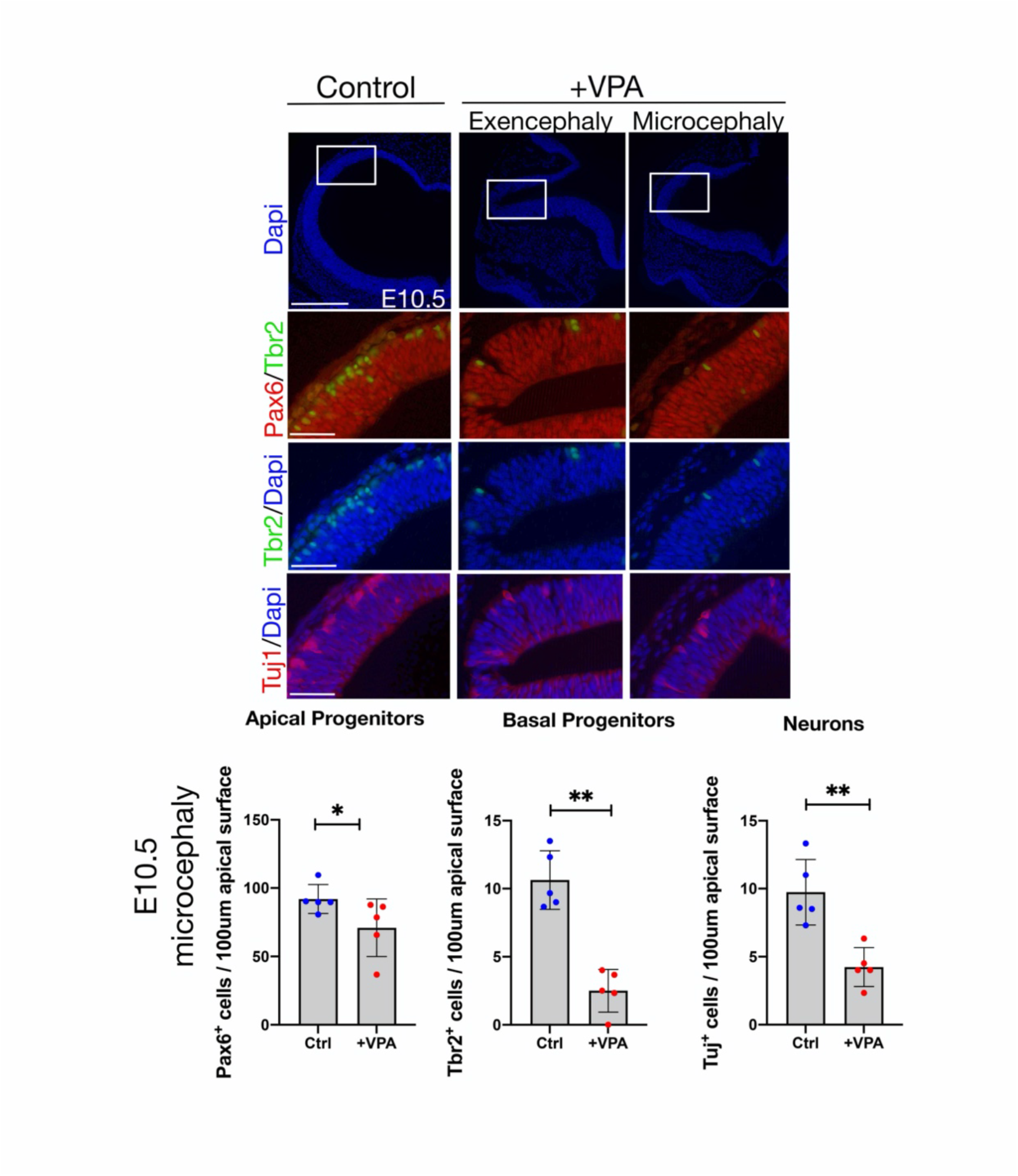
Impaired neurogenesis after VPA treatment is already observed at E10.5. Cortical sections (coronal) of E10.5 embryos were immunostained for Pax6, Tbr2, Tuj1 and counterstained with Dapi. Scale bar, 250μm (top row), 50μm. Graphs show quantification of Pax6 and Tbr2 positive progenitors or the thickness of the neuronal layer in the microcephalic cortical vesicles (5 embryos from at least 4 different mothers were analyzed). Data bars represent mean ± SEM Mann-Whitney test: *p ≤ 0.05 and **p ≤ 0.01.

**Figure S3.**
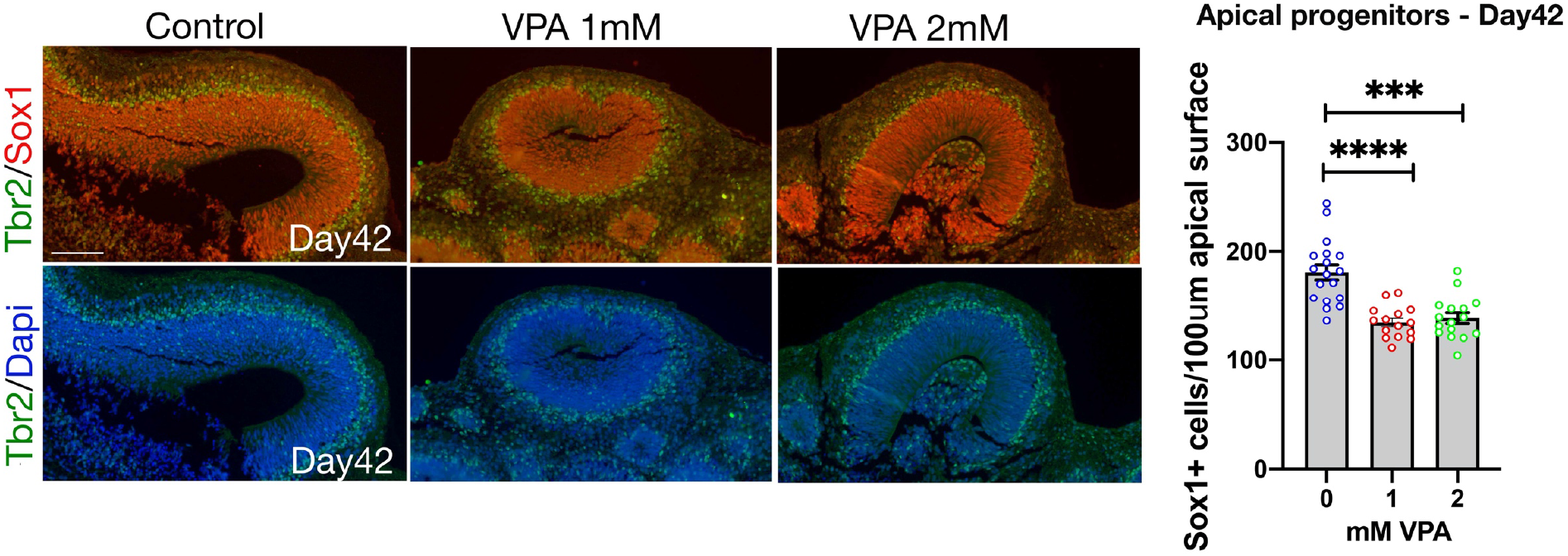
Neurogenesis is impaired in cerebral organoids treated with VPA. Sections through control and VPA-treated organoids were immunostained with Sox1(red), Tbr2 (green) and Dapi (blue) at day 42 (Scale bar, 50μm), (*n* = 15 (Control), 12 (1mM VPA), 13 (2mM VPA), 4 independent experiments). Kruskal-Wallis test: ****p ≤ 0.001 and ****p ≤ 0.0001.

**Figure S4.**
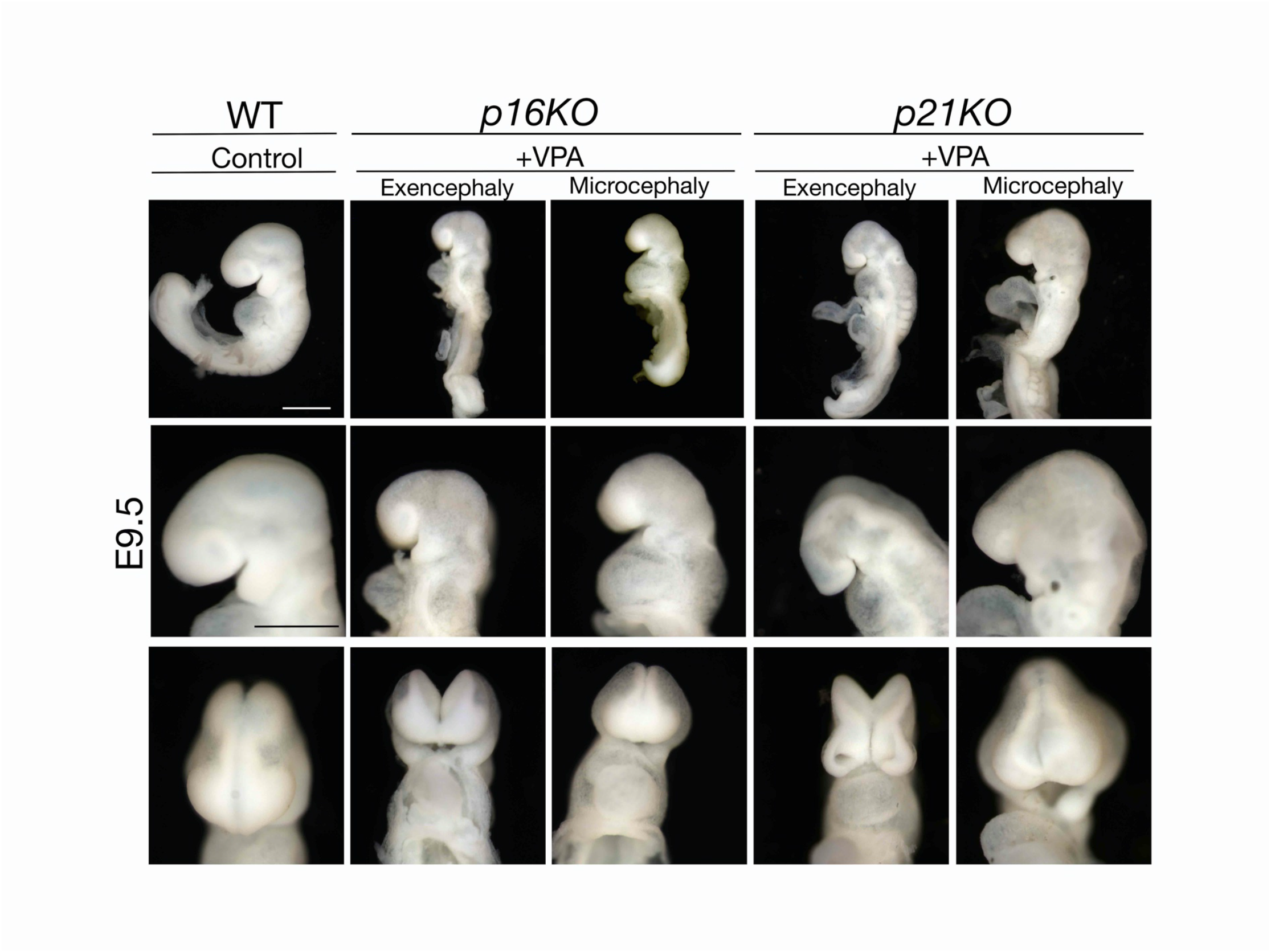
Genetic deficiency of *p16^Ink4a^* or *p21* does not rescue VPA induced phenotypes. Lateral views of control and VPA-treated embryos deficient for *p16^Ink4a^* or *p21* (top row). Scale bar, 500μm. Higher magnification of the heads in lateral (middle row) and frontal views (bottom row). Scale bar, 50μm. An open neural tube or a smaller brain, as well as a gross misalignment of the neural tube and somites are still observed after VPA treatment in the absence of *p16^Ink4a^* or *p21*.

**Figure S5.**
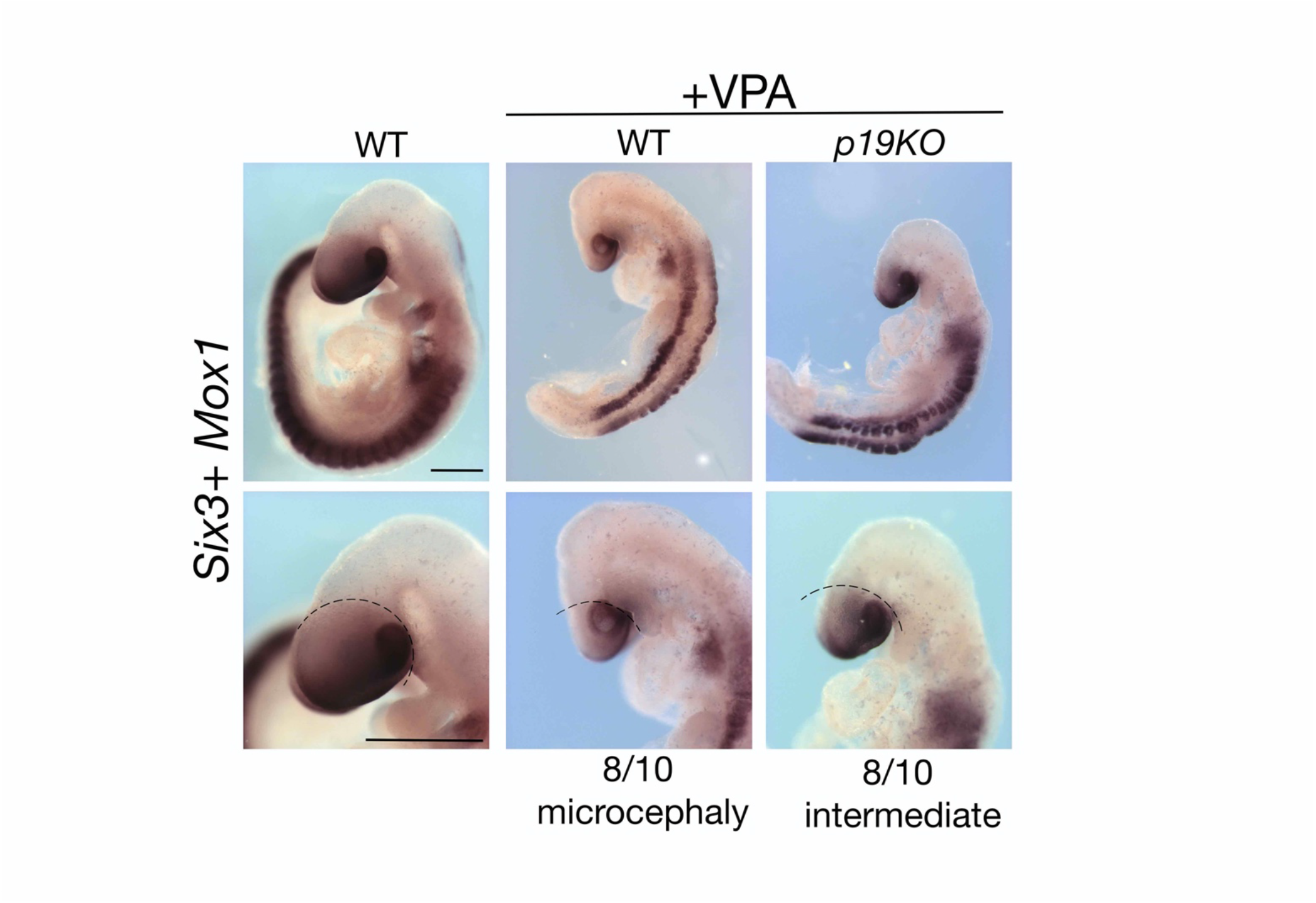
Improved forebrain phenotype in *p19^Arf^* deficient mice after VPA treatment. Whole mount *in situ* hybridization for *Six3* (forebrain) and *Mox1* (somites), showing an increased size of the forebrain in *p19^Arf^*-deficient, VPA-treated mice in comparison to the *WT* mice treated with VPA. Scale bar, 500μm (top row) and 50μm (bottom row). The number of embryos examined are indicated (*n*=10 from at least 5 different litters).

**Figure S6.**
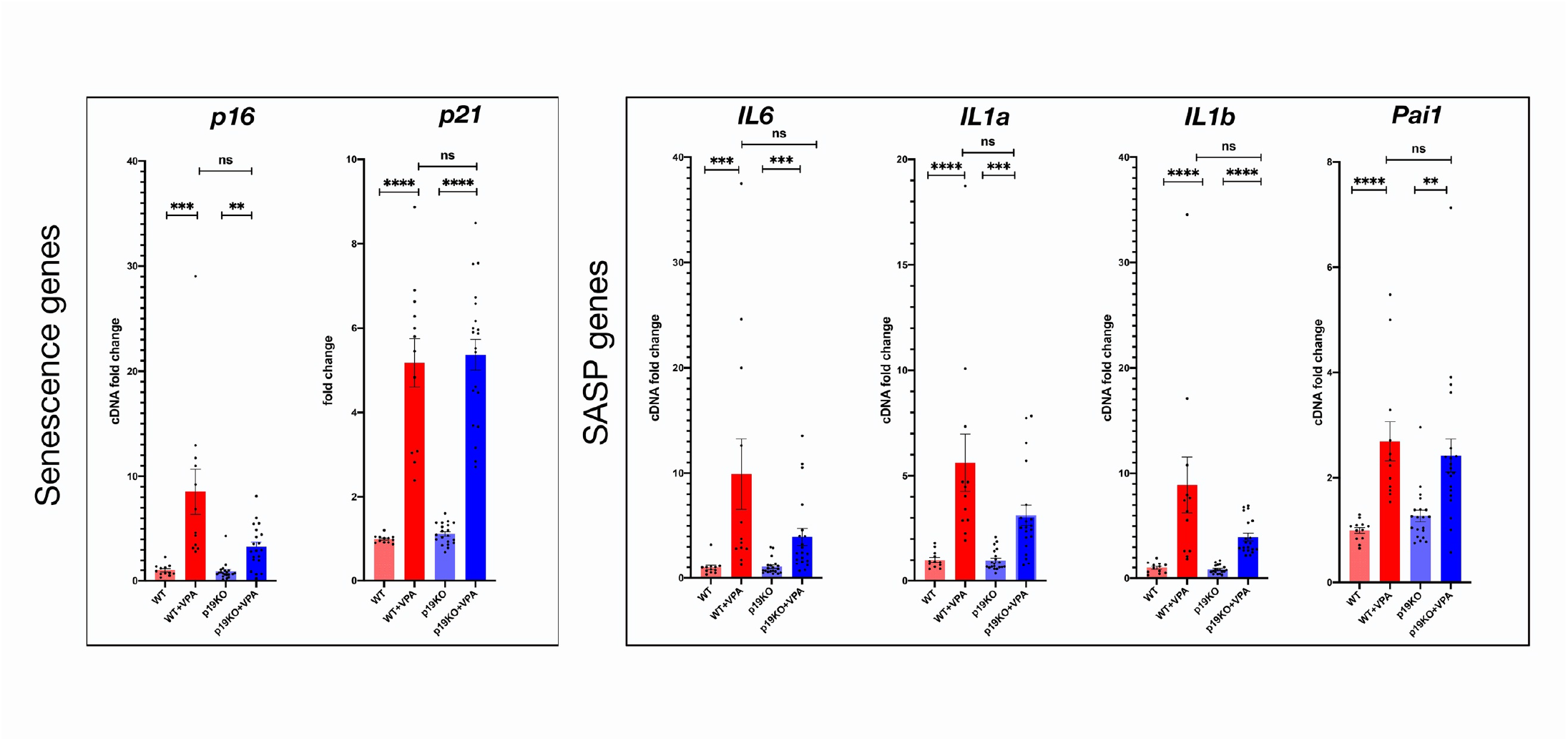
Senescence and SASP genes are less induced in *p19^Arf^*-deficient embryos with VPA treatment. qRT-PCR analysis on E8.75 forebrain and midbrain, from control and *p19^Arf^*-deficient mice, treated with VPA or left untreated. Graphs show fold change expression for the senescence markers (*p21, p16^Ink4a^*) and for SASP genes (*IL6, IL1a, IL1b*, and *Pai1*), normalized to untreated control (*n* = 12 (Control), *n*=12 (Control+VPA), *n*=20 (*p19KO*), *n*=20 (*p19KO*+VPA), from at least 3 different litters). Data bars represent mean ± SEM. Kruskal-Wallis test: ns, no significant, **p ≤ 0.01, ***p ≤ 0.001 and ****p ≤ 0.0001.

**Figure S7.**
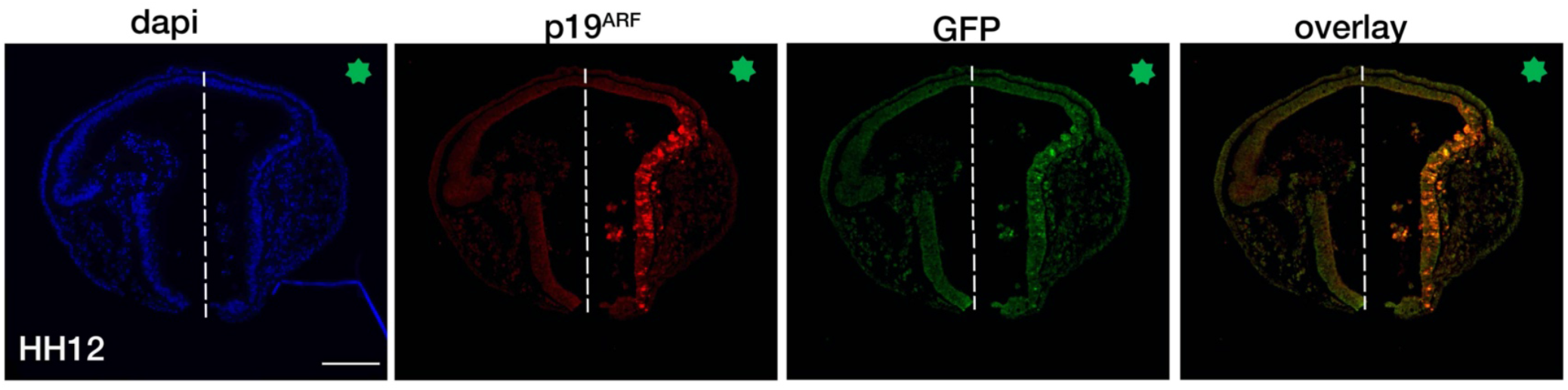
Ectopic expression of *p19^Arf^ -GFP* in the chicken neural tube. Sections through the neural tube of *p19^Arf^-GFP* electroporated chicken embryos electroporated at stage HH12, immunostained for p19^Arf^ (red) and GFP (green), with Dapi counterstaining (blue). The green star shows the electroporated side. Scale bar, 100μm.

**Figure S8.**
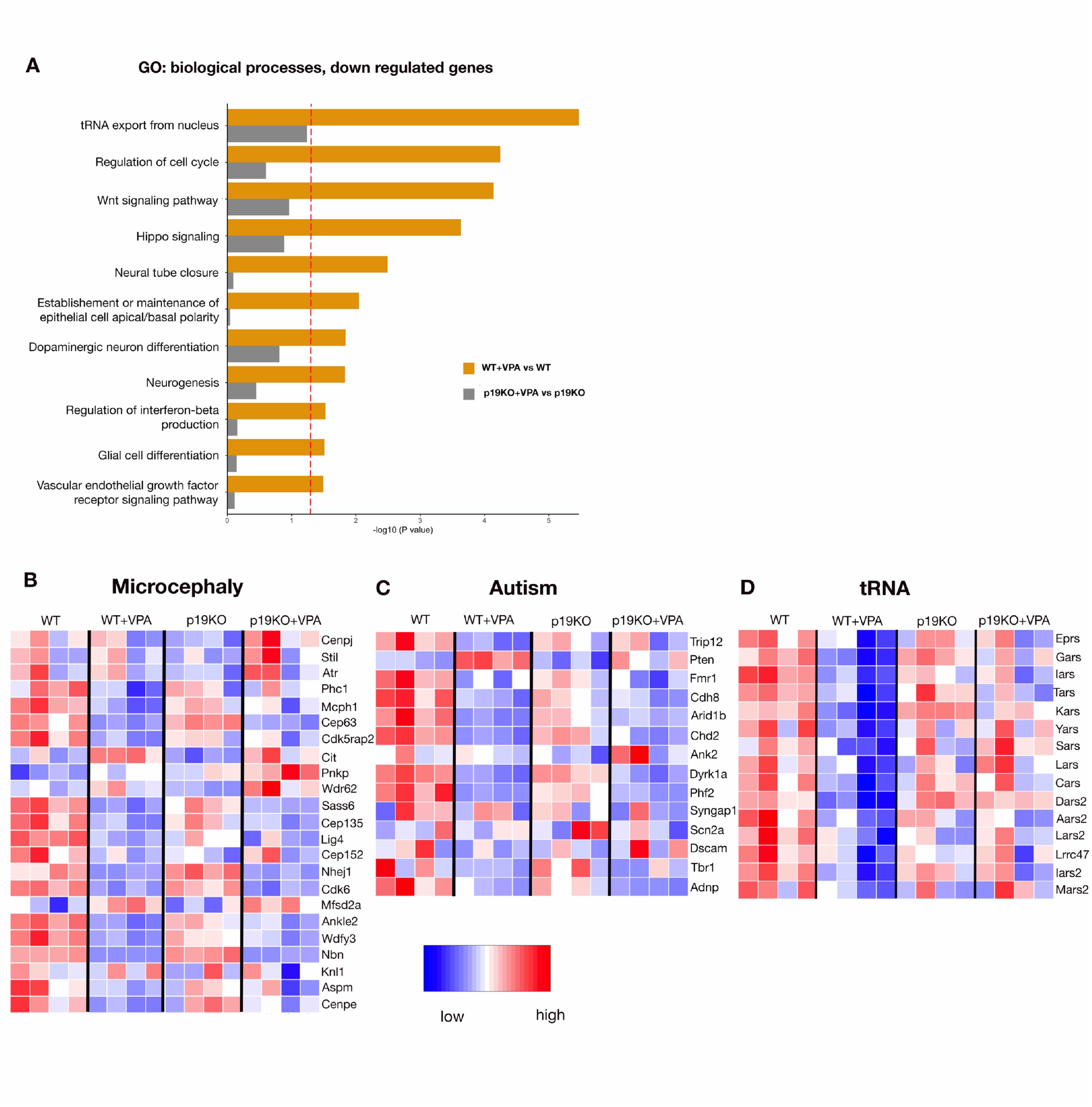
RNA-sequencing data analysis uncovers neurodevelopmental and tRNA related signatures as being less affected by VPA treatment in *p19^Arf^*-deficient mice. **(A)** GO Biological Processes pathway analysis on the downregulated genes from RNA-seq of the forebrain and midbrain. Heat Maps showing the relative expression of representative genes associated with **(B)** Microcephaly (list generated from Jayaraman et al, 2018), **(C)** Autism (List generated from Jayaraman et al, 2018) and **(D)** tRNA (list of genes identified in Fig. 6C pathway analysis).

**Table S1.**
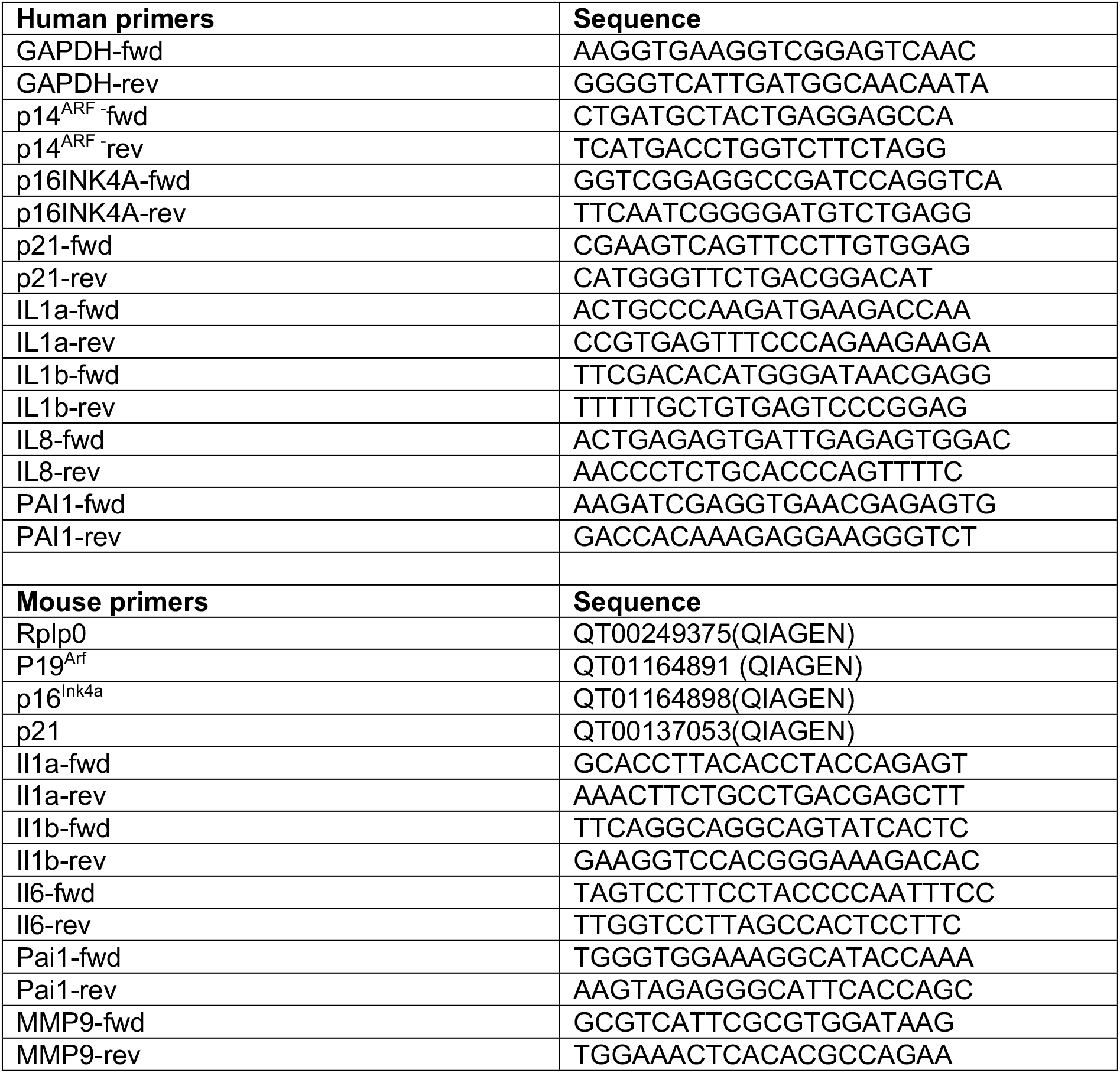
Primers used for qRT-PCR in the study. The table lists the primer sequences or source for both human and mouse genes used in the study.

